# S’Wipe: User-Friendly Stool Collection for High-Throughput Gut Metabolomics

**DOI:** 10.1101/2025.02.27.640581

**Authors:** Dana Moradi, Ali Lotfi, Alexey V. Melnik, Konstantin Pobozhev, Hannah Monahan, Evguenia Kopylova, Yanjiao Zhou, Alexander A. Aksenov

## Abstract

Microbiome is increasingly recognized as a key factor in health. Intestinal microbiota modulates gut homeostasis via a range of diverse metabolites. For example, molecules such as short chain fatty acids (SCFAs), the microbial fermentation products of dietary fiber, have been established to be reflective of microbiome and/or dietary shifts and SCFAs alterations have been linked to multiple gastrointestinal disorders from cancer to colitis. Despite their potential as biomarkers, technical challenges in stool collection have limited clinical translation. Here we present Stool Wipe (S’Wipe), an ultra-low-cost fecal collection method using lint-free, mass spectrometry-compatible cellulose wipes as toilet paper. Specimens are preserved in ethanol without refrigeration and can be shipped via regular mail. Mass spectrometry analysis demonstrated that S’Wipe captures both volatile and non-volatile metabolites with reproducibility and stability validated for diagnostically relevant molecules. We show that S’Wipe performs equivalently to direct stool collection, enabling interchangeable use and comparison with existing studies. This methodology is ideally suited for large-scale population studies, longitudinal tracking, and personalized medicine applications.

**IMPORTANCE:** Gut microbiome and intestinal metabolome present invaluable diagnostic and therapeutic targets. However, conventional stool testing has several barriers limiting bioassessment from populations. Routine, high temporal resolution monitoring of stool metabolome, including extensively validated and broadly informative biomarkers such as short chain fatty acids (SCFAs), is not implemented due to relatively high cost and inconvenience of sampling, possible need for clinical setting for sample collection, difficulty to collect samples reproducibly, especially due to potential for user errors, requirement for freezer storage and maintaining cold chain during shipment. We present a sampling strategy specifically designed to overcome these obstacles. We demonstrate how this method can enable capturing accurate molecular snapshots at massive scales, at ultra low cost. The approach collapses complex medical-grade collection into easy self-administration. Individuals can thereby self-monitor therapeutic responses through routine metabolome tracking, including the volatilome, otherwise hindered by infrastructure restrictions. Ultimately, this sampling approach is intended to enable participatory wellness transformation through practical high frequency self-sampling.

## Introduction

The gut microbiome plays a central role in human health in profound and wide-ranging ways through nutrient metabolism, immune modulation, and metabolic regulation, including production of bioactive metabolites (1–3). From numerous investigations of gut-derived molecules, short-chain fatty acids (SCFAs) have emerged as particularly important biomarkers of intestinal health. These bacterial metabolites have been extensively validated as key gut health indicators, with studies demonstrating their influence on processes spanning digestion, metabolism, immunity, and even mental health (4–6). SCFAs can be measured across the gut, with fecal samples being the most accessible (7–10). SCFAs are predominantly produced through fermentation of dietary fibers, subsequently permeate throughout the body, and affect a diverse range of tissues and physiological processes (10–12). The clinical potential of these microbial metabolites as measurable biomarkers continues to be unraveled across investigations - from IBD, all the way to the inflammatory cascade and metabolic syndrome underpinning contemporaneous epidemics of obesity and cardiovascular disease afflicting aging populations (13–17). SCFAs deficiencies or disbalances have been associated with low-fiber intake diets, altered gut permeability and inflammation, thereby promoting insulin resistance, elevating risk factors, and increasing the probability of cardiovascular events (18–21). SCFAs quantification and profiling offer clinical opportunities for non-invasive monitoring, not only for disease diagnostics, but potentially early risk assessment and exposure-response monitoring, as well as guidance for nutritional or therapeutic gut microbiome modulation. Still, SCFAs represent just one class of informative gut metabolites. Other molecular markers detectable in stool include bile acids, which reflect lipid metabolism and gut barrier function; p-cresol and indole, which indicate protein metabolism and are linked to various health conditions, such as Autism Spectrum Disorder (ASD)(22); and a range of vitamins, amino acids, and lipids that provide insights into both dietary habits and microbial activities. Together, these diverse molecular families create a comprehensive window into intestinal health status.

Molecular patterns in the gut microbiome, while linked to various health conditions, remain challenging to translate into clinical biomarkers due to high biological and technical variability. While research has established clear connections between gut metabolites and health, the clinical implementation requires robust reference ranges and validated associations with health outcomes. These can only be established through extensive longitudinal sampling to separate consistent molecular signatures from background variability. However, current sampling methods present significant practical barriers to such large-scale data collection. Fecal samples rapidly degrade in ambient conditions due to chemical degradation and microbial growth, while reliance on specialized collection and handling infrastructure, cost, and complexity of the collection, coupled to a wide range of non-standardized sampling protocols poses various logistical challenges. Consequently, at the present time, both patients and the general public do not have access to reliable personalized monitoring of gut molecular biomarkers. Last, but not the least, collecting feces is objectionable to many people, limiting their willingness to participate in studies that require fecal testing, especially those that entail frequent, e.g. longitudinal collection.

Current stool sampling collection methodologies for fecal sampling for microbiome and metabolomics analysis used in clinical practice are predominantly traditional stool collection kits consisting of stool hats, swabs, or small containers (23–29). Typically, stool needs to be collected in a screen and then manually transferred into a storage container (e.g. Cologuard® test). For some methods, samples must be immediately frozen and then transported frozen to a lab to help preserve their integrity. This can lead to high variability due to differences in storage and transportation temperature, transit time and exact collection methods between patients. Another example of the currently used method is fecal occult blood test (FOBT) cards (30–34). In FOBT sampling fecal smears are collected at home on specialized sample cards designed to detect blood. Those cards help stabilize some molecules for transport at room temperature. However, drying alters the molecular composition. Also, the small sample size and difficulties in using and handling cards limit broader profiling. Specialized solutions such as the Cologuard® can be used for non-invasive tests that analyze stool samples patients collect on a commercial kit with a preservative buffer. This test uses genetic sequencing to detect colorectal precancers but is not optimized for metabolomics (33–35). For the majority of stool-based tests (screening for blood in the stool, stool culturing for bacterial or viral infections diagnostics, ova and parasite exam, fat malabsorption assessment, etc.), stool is collected in research labs or hospitals, where clinicians may collect fecal samples directly, or during procedures like lavage or endoscopic biopsies. Obviously, such invasive techniques limit practical scalability (33).

Overall, sampling approaches that are scalable, patient- and consumer-friendly and preserve integrity during non-refrigerated transportation, have remained an unmet need. Most testing today still relies on traditional immediate frozen collection protocols (23, 25), which are logistically impractical for population-level studies. To bridge this gap, we developed the S’Wipe, a user-friendly, ultra low-cost fecal collection method. A cellulose collection paper is used as lavatory towels, which makes sampling identical to a regular bathroom routine. Using GC-MS and LC-MS, we demonstrate that S’Wipe captures native SCFAs and other diverse diet-/host-associated metabolites at highly reproducible levels over weeks, without cold storage. The DNA is captured on the cellulose and can be used for complementary sequencing. The S’Wipe method is designed to enable democratized, scalable gut molecular monitoring to stimulate personalized solutions across global populations.

## RESULTS

### S’Wipe

The Stool Wipe (S’Wipe) sampling approach consists of three key components: (1) a lint-free cellulose collection paper that serves as the sampling matrix, (2) a collection tube containing preservation solution (60% ethanol is utilized throughout this study), and (3) a standardized protocol for sample handling and processing. The name ‘S’Wipe’ refers to this complete sampling system, where the collection paper is used like conventional toilet paper during normal bathroom routines, then placed in the preservation solution for storage and shipping.

With S’Wipe use, approximately 100 mg of stool can be typically collected. The wipe is then placed by the user into a 20 mL wide-mouth centrifuge tube with 5mL of MS-grade 60% ethanol spiked with internal standard, sealed, and then can be both stored and/or shipped at room temperature until extraction and analysis (Figure 1). The users are not required to modify their bathroom routine with the exception of placing the used paper into a collection tube rather than flushing/throwing it away. Due to the simplicity of collection, sampling can be conducted anywhere, and every bowel movement could be sampled, if needed. Correspondingly, large longitudinal sampling with each subject being their own control for intervention studies can be facilitated. During shipment, the extraction solvent remains absorbed by the wipe and acts as a preservative. Upon arrival at the analysis lab, the samples are centrifuged to separate the supernatant from the wipe. The supernatant is directly compatible with mass spectrometry analysis, both GC-MS and LC-MS, without any additional sample preparation steps, with the exception of possible dilution. The absolute measured abundances of biomarkers of interest, as well as longitudinal trends can then be used to inform users.

**Figure 1.**
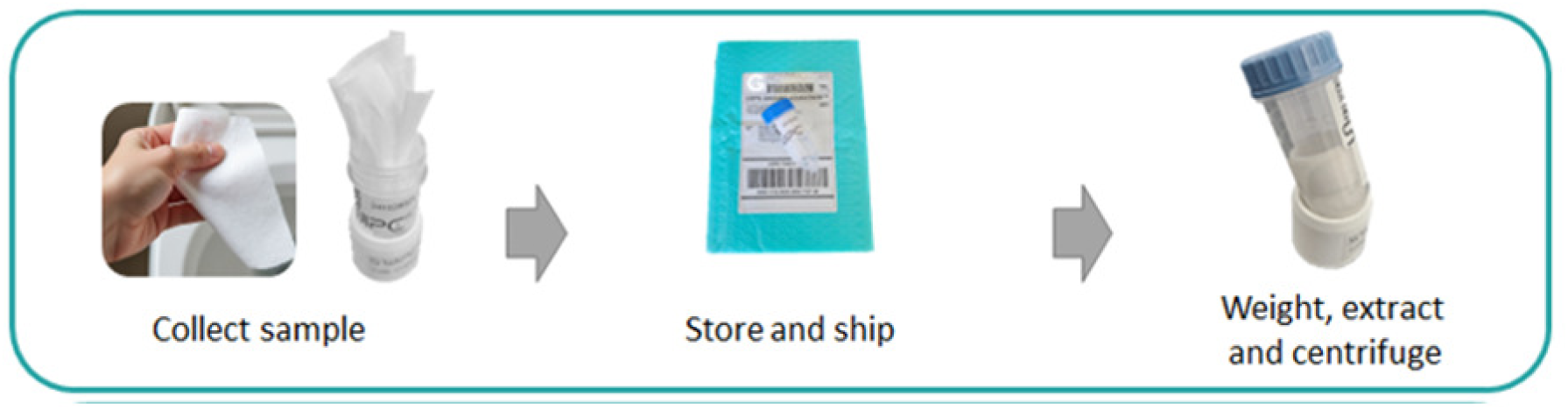
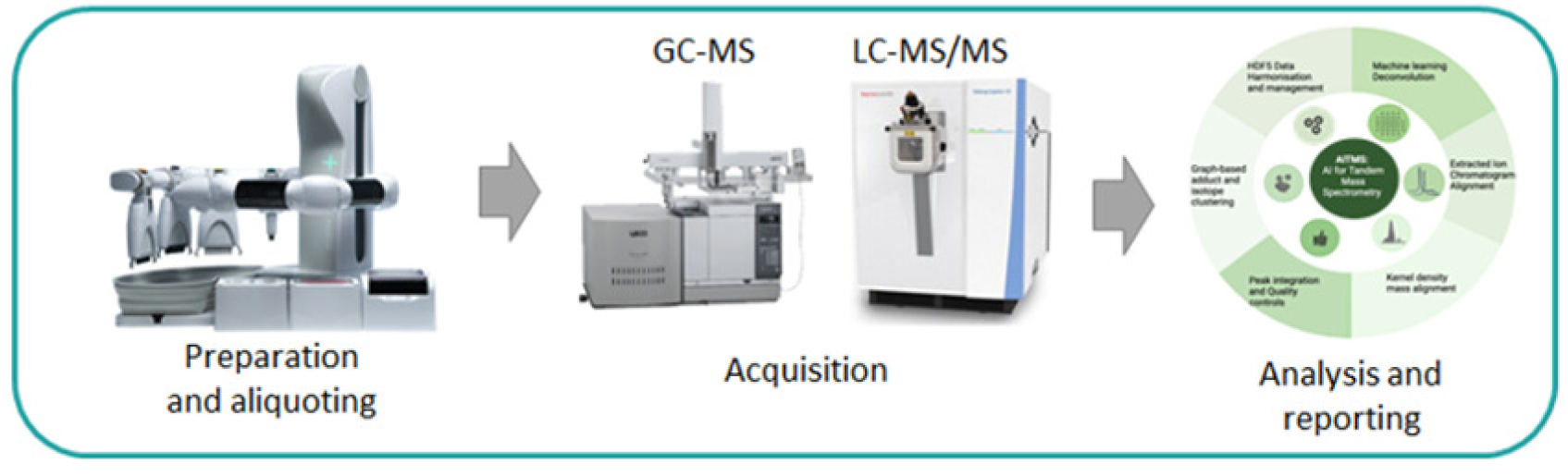
S’Wipe* sampling. **A)** Sampling and shipping workflow using the kit (sample collection kit is shown in Supplementary Figure S1 (10.6084/m9.figshare.28404761, https://figshare.com/articles/figure/Figure_S1_/28404761?file=52313852). **B)** Schematic sample preparation and analysis workflow. *The patent application is pending (US Application #63/570,322).

### Sampling

The S’Wipe sampling was found to capture compounds of interest such as SCFAs, p-cresol, and other molecules that are expected to be detectable in stool, such as indole, bile acids, vitamins, amino acids, lipids, etc. (see Methods section, Protocol Optimization study, also Supplementary Figure S2 (10.6084/m9.figshare.28404800, https://figshare.com/articles/figure/Figure_S2_/28404800), S4 (10.6084/m9.figshare.28404827, https://figshare.com/articles/figure/Figure_S4/28404827) and S5 (10.6084/m9.figshare.28404872, https://figshare.com/articles/figure/Figure_S5/28404872)). While molecular networking demonstrates that S’Wipe preserves the broader stool metabolome, we specifically focused our validation on SCFAs and other established diagnostic markers to rigorously assess the method’s performance for molecules of immediate clinical relevance. As S’Wipe allows for measuring the stool weight, the abundances of metabolites could be normalized (Figure 2A; see Methods section, Stability study). Also, with the use of external calibration, absolute quantities for various molecules could be calculated and used as biomarkers to gain lifestyle or other information, e.g. Supplementary Figure S2A,B (10.6084/m9.figshare.28404800, https://figshare.com/articles/figure/Figure_S2_/28404800). (see Methods, Interpersonal Variability Study) The ratios of metabolites are not dependent on the absolute quantitation, and can also be more linked to specific conditions (36–38)-(39–42), (43) (Supplementary Figure S2D (10.6084/m9.figshare.28404800, https://figshare.com/articles/figure/Figure_S2_/28404800)).

**Figure 2.**
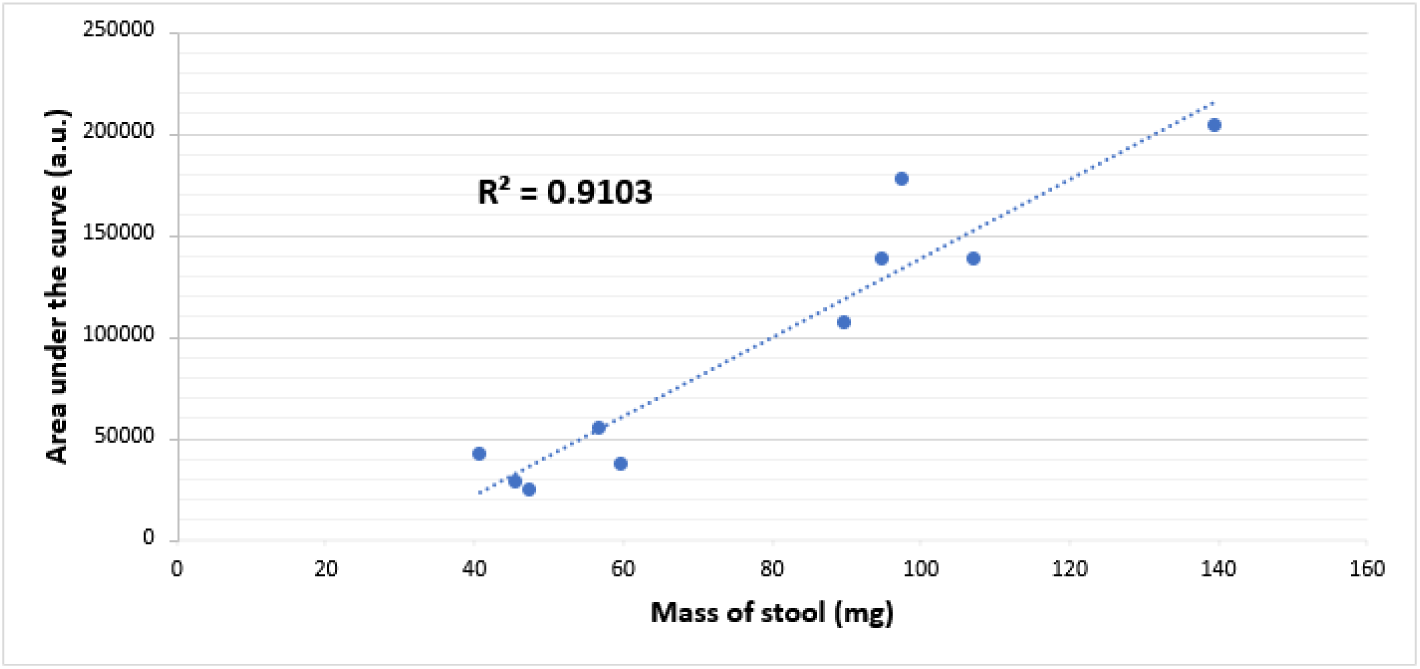
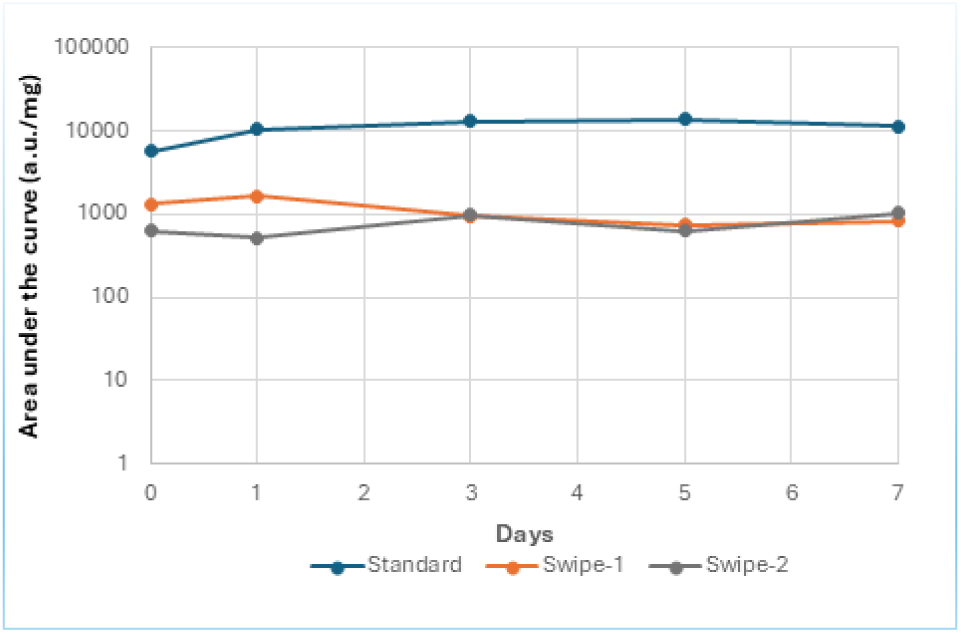
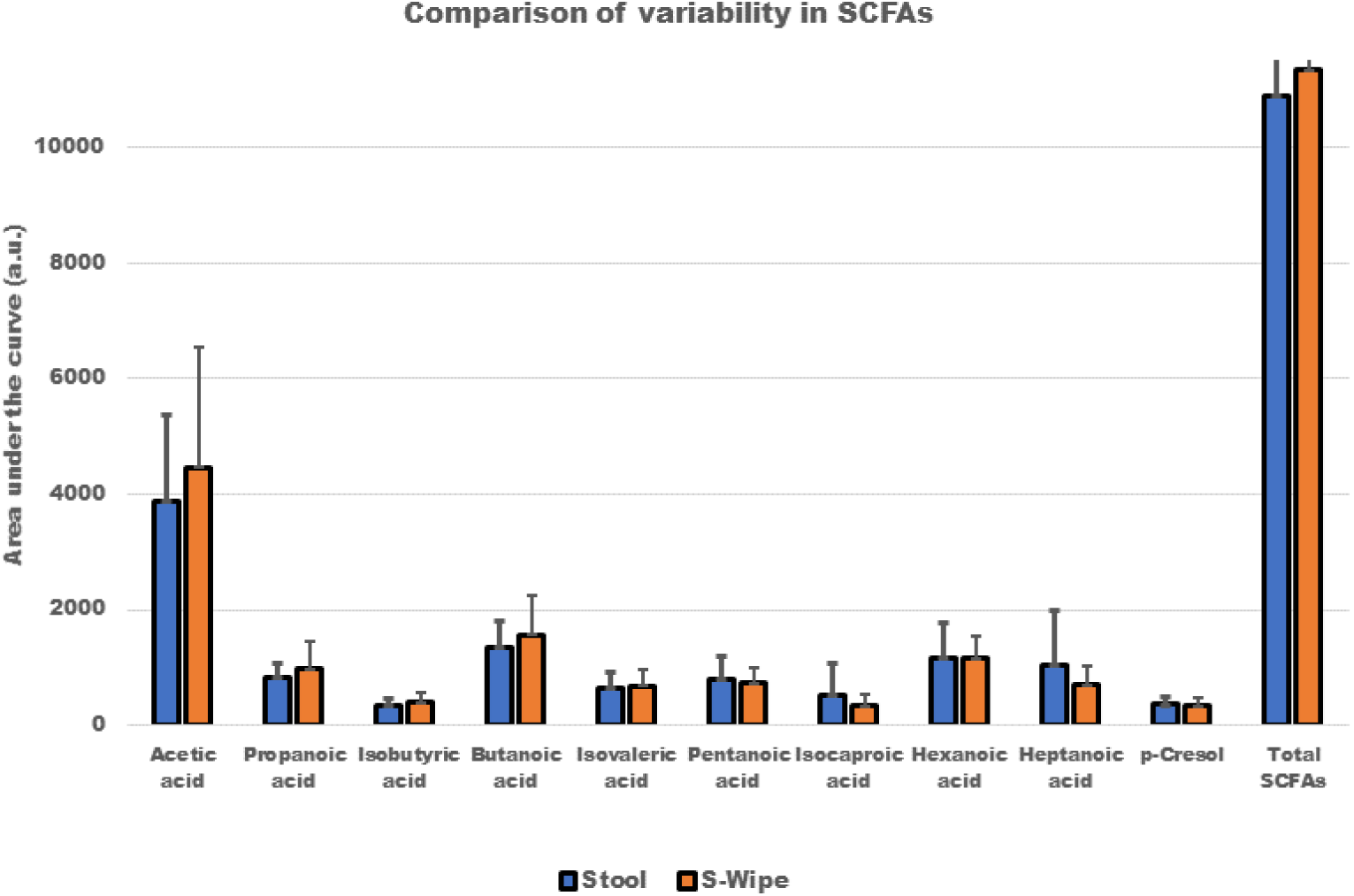
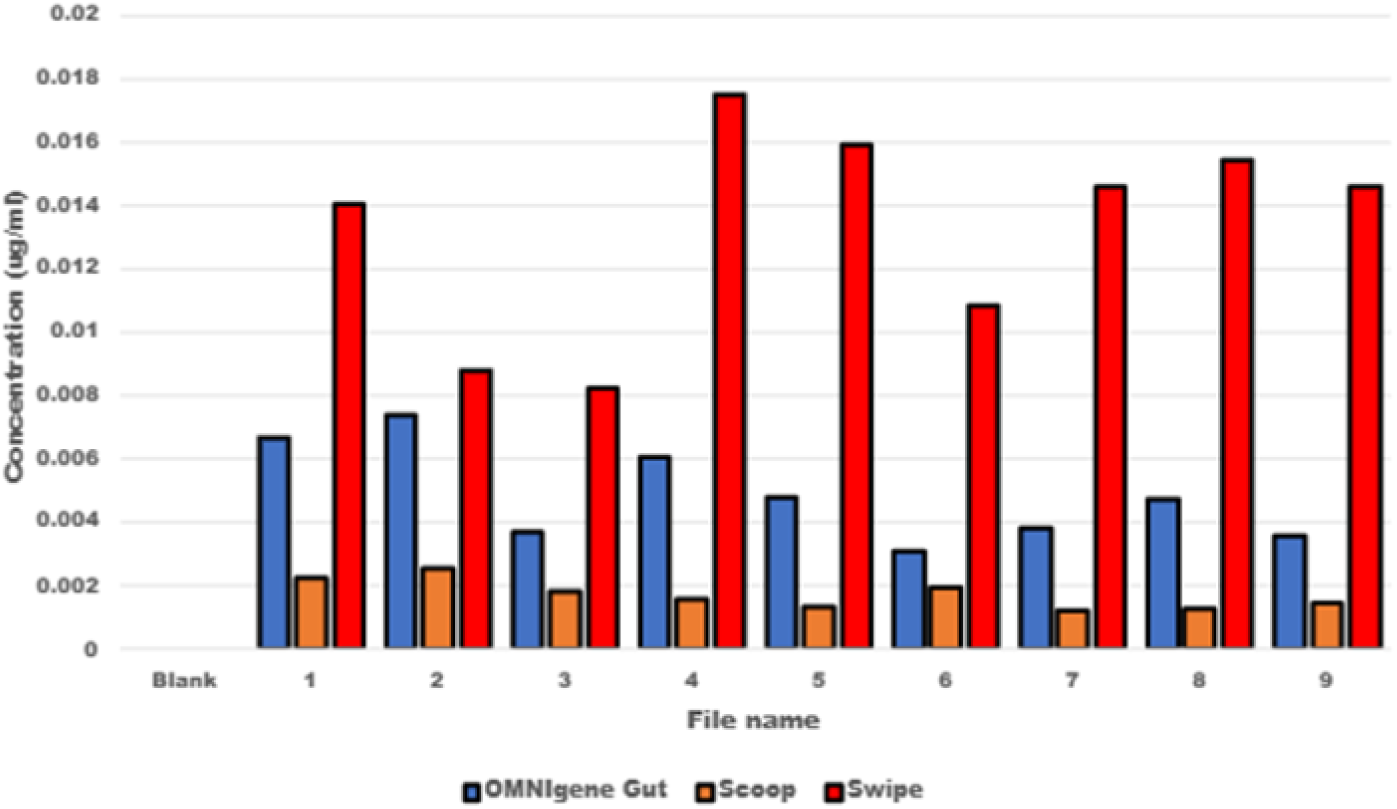
Methodology Benchmarking Comparison. **A)** Scatter plot of total abundance of C2-C7 SCFAs vs. the measured mass of collected samples for both stool and S’Wipe. A significant positive correlation (Pearson’s R = 0.95, p < 0.001)) is evident. **B)** Stability of SCFAs**. C)** Comparison of variability in SCFAs for stool collected with scooping method (conventional sample collection) vs. S’Wipe collection. **D)** Plot showing the total concentration of SCFAs in supernatant for different users, normalized by the stool weight. Scoop - regular stool collection (Blue); OMNIgene Gut (Orange) and S’Wipe (Red).

### Stability during storage

To evaluate S’Wipe’s performance for SCFA analysis, we first assessed the stability of these metabolites under room temperature storage conditions. While SCFAs are generally stable at room temperature, their preservation specifically within the S’Wipe matrix needs to be validated. We conducted two complementary experiments: first, we spiked S’Wipes with a 500μl aliquot of Accustandard FAMQ-004 standard (New Haven, CT USA) at 10mM concentration to track recovery of known SCFA concentrations. Second, we collected real stool samples using S’Wipe to assess the stability of endogenous SCFAs. Both the standard-spiked wipes and stool-sampled wipes were stored at room temperature for periods ranging from one day to one week.

GC-MS analysis revealed notable stability of all nine SCFAs over the study period. Statistical analysis showed an average relative standard deviation of 23.48% across all time points (see Methods, Stability study), demonstrating good technical reproducibility. Most importantly, no noticeable degradation was observed during room temperature storage (Figure 2B, Supplementary Figure S6A (10.6084/m9.figshare.28404884, https://figshare.com/articles/figure/Figure_S6/28404884)).

### Benchmarking

To benchmark S’Wipe against existing methods, we conducted side-by-side comparisons with conventional stool collection and commercial sampling approaches. Previous studies comparing preservation methods (44) demonstrated that 95% ethanol provides excellent metabolite preservation compared to FOBT and FIT cards, though high ethanol concentration can complicate sample handling and processing. Building on these findings, we compared S’Wipe using 60% ethanol, with direct collection and OMNIgeneene Gut. The latter is chosen as a current commercially available standard for room-temperature metabolomics sampling. The 60% ethanol concentration is chosen to balance preservation with practical handling considerations (particularly reduced flammability), while maintaining similar metabolite recovery to direct collection methods, as demonstrated in Figure 2. We focused particularly on three SCFAs with established diagnostic importance - acetic acid, propanoic acid, and butyric acid - while also monitoring other SCFAs and p-cresol (Figure 2C, Supplemental Table1, Supplemental Figure S6 (10.6084/m9.figshare.28404884, https://figshare.com/articles/figure/Figure_S6/28404884), Supplemental Figure S7 (10.6084/m9.figshare.28404908, https://figshare.com/articles/figure/Figure_S7/28404908), Supplemental Table 2). The comparison revealed that S’Wipe captured these metabolites with comparable or better sample-to-sample consistency than traditional methods (Figure 2D and Supplemental Figure S8 (10.6084/m9.figshare.28404911, https://figshare.com/articles/figure/Figure_S8/28404911?file=52314176)). While measured concentrations aligned well across all three collection methods, S’Wipe demonstrated lower variability in replicate measurements. One notable exception was acetic acid, which showed high variability across all methods, ostensibly due to its high volatility.

In the proposed approach, ethanol solution is used for sample preservation. The ethanol is known to both denature enzymes, preventing enzymatic degradation (45–49), and kill microorganisms preventing blooms (45, 49–51). Focusing on the SCFAs, it is evident that S’Wipe-collected organic acid remained stable for at least several days at ambient temperature. GC-MS analysis indicated negligible changes, with no statistically significant degradation trends over the entire time interval (p > 0.24) (Figure 2B). Quantitative linearity of short-chain fatty acid detection spanned physiologic concentrations (Supplementary Figure S3 (10.6084/m9.figshare.28404812, https://figshare.com/articles/figure/Figure_S3_/28404812)). Absolute concentrations aligned with values reported from conventionally collected samples, confirming compositional integrity.

### Stability during shipping

The intended use of S’Wipe assumes at-home self-sampling, followed by shipping via regular mail. To assess the stability of metabolome under typical shipping scenarios, we have further conducted a shipping/storage experiment, where the samples were compared between different handling conditions: refrigerated storage, room temperature storage, short- and long-distance shipping (Figure 3B). For each handling condition we had three replicants. Correspondingly, the samples collected by S’Wipe were stored at −80C immediately after collection, stored at room temperature for the duration of the study (one week), shipped, then returned, locally (within the state) and shipped/returned long distance (from the East to West coast of the US). Supplemental Table 3 shows the Standard deviation for different SCFAs under each sample handling condition. The shipping reflects a representative sum total of fluctuations in ambient conditions, in particular exposures to high temperatures and humidity. The measured abundances of SCFAs are consistent across all three sample handling scenarios, indicating that their degradation did not occur noticeably (Figure 3B). Consequently, such a handling strategy is at least appropriate within the outlined conditions for the molecules of interest described here.

**Figure 3.**
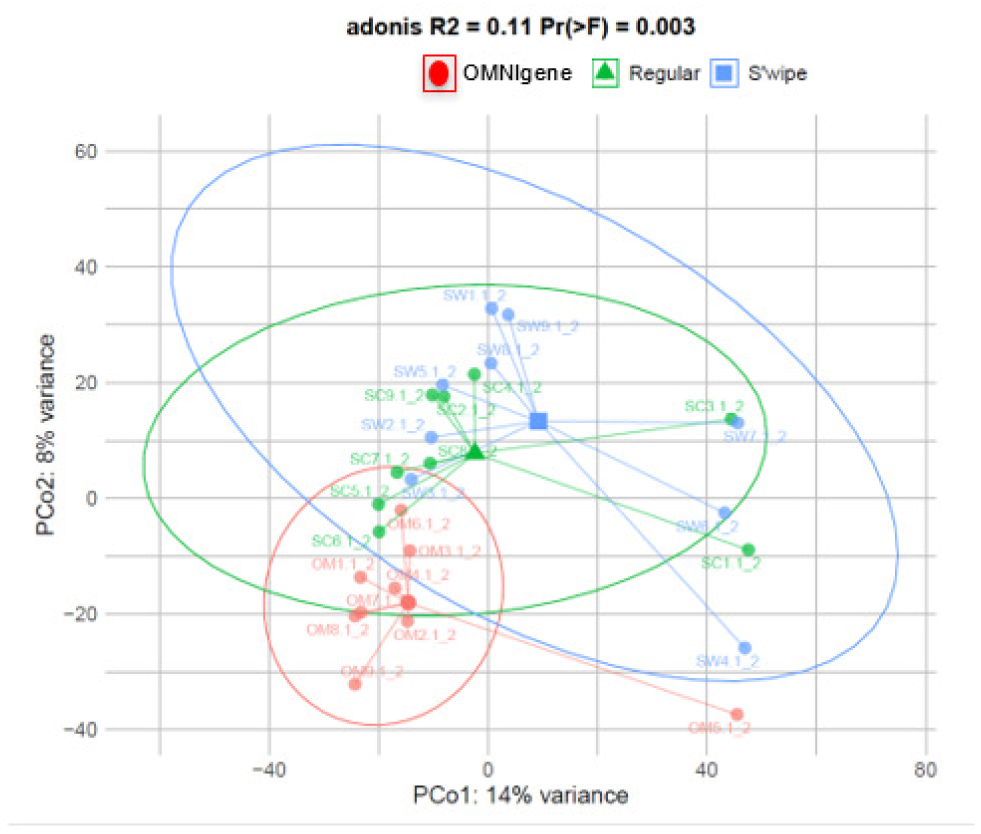
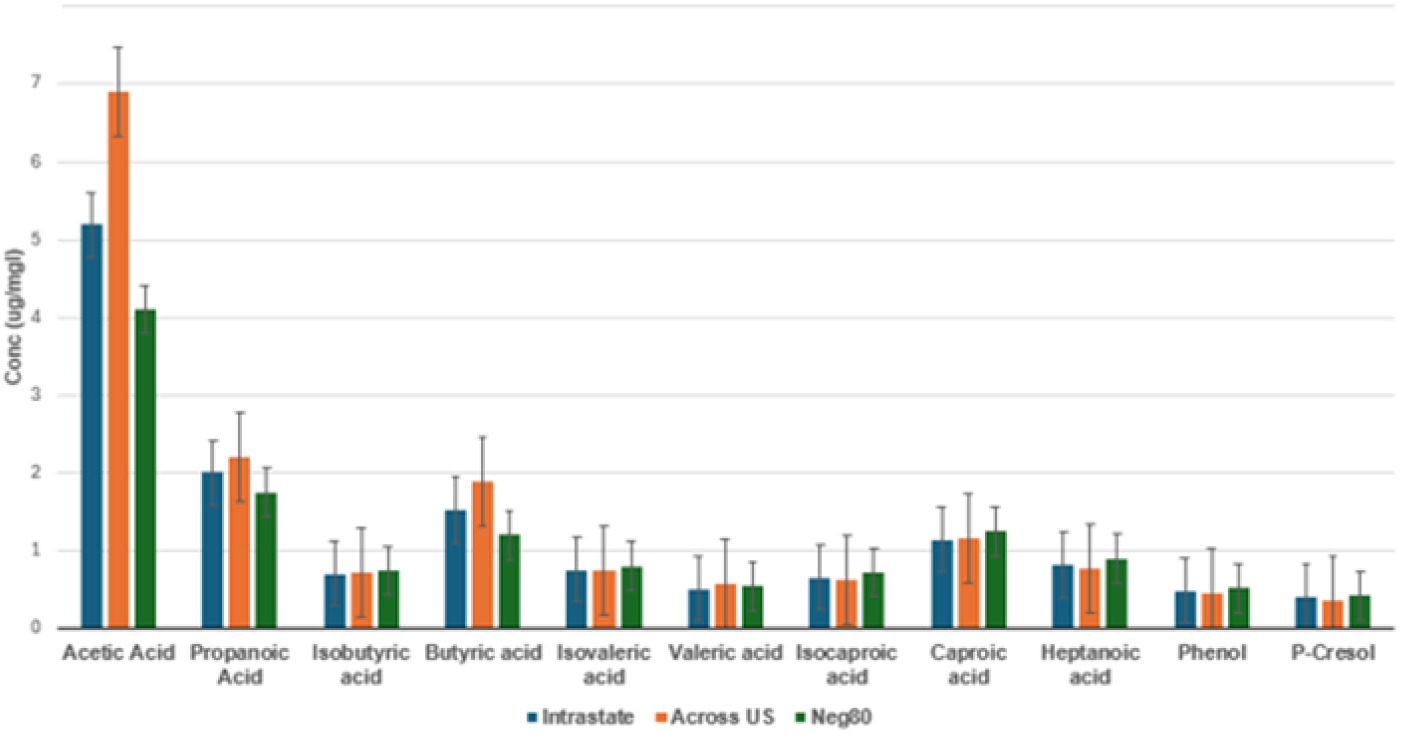
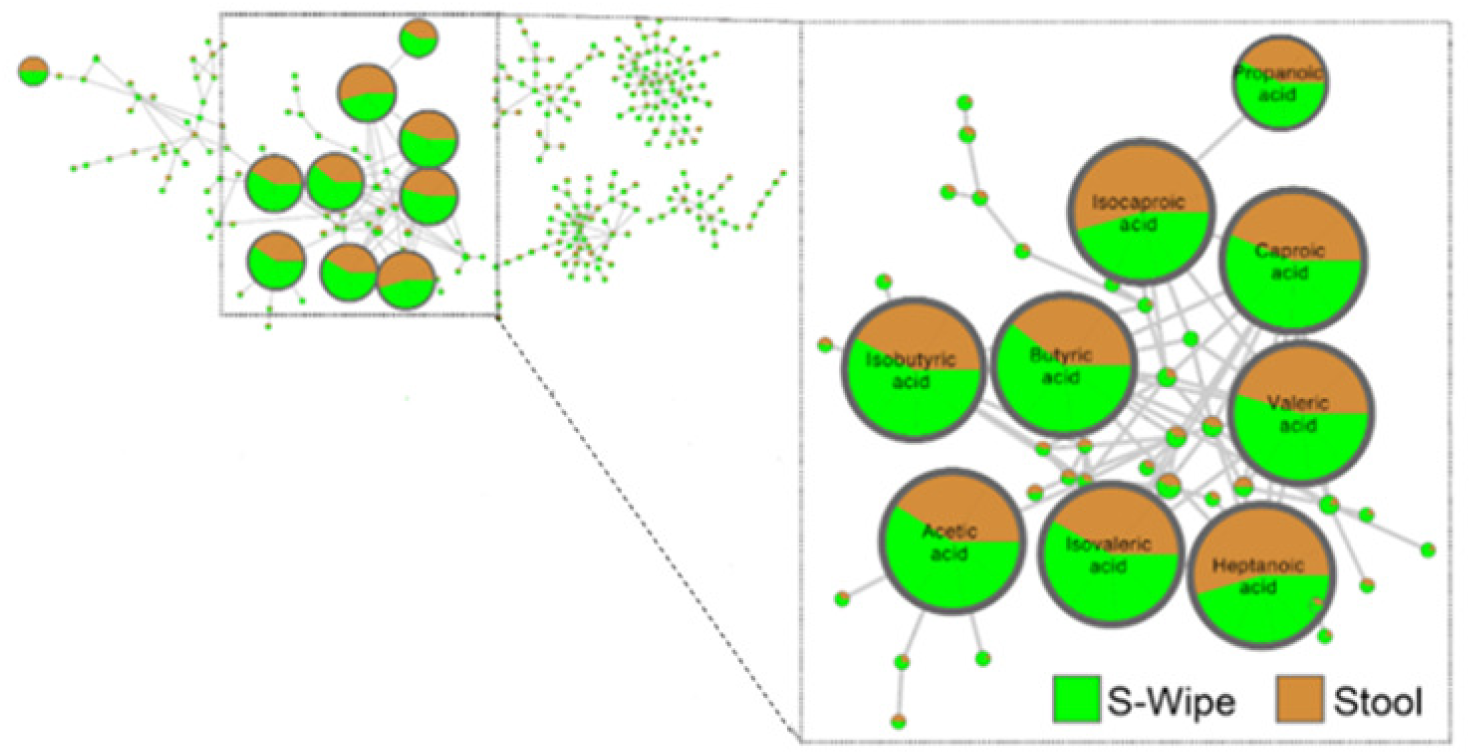
Stability study. **A)** Principal coordinate analysis plot of samples from the conventional stool collection and S’Wipe. Several extraction protocols have been tested, as described in the Methods section, to account for possible biases due to the sample preparation. **B)** Bar plot showing the distribution of short- and medium-chain fatty acids as well as p-cresol and phenol across three different conditions in the shipping study. Neg80: samples stored at −80°C immediately after collection. Intrastate: samples shipped within the state (approximately one week passed in between shipping and receiving samples). Across US: samples shipped from East to West coast of the US (approximately two weeks passed in between shipping and receiving samples). The “Intrastate” and “Across US” samples were stored at 2°C-4°C upon arrival to the lab until analysis. All of the samples were analyzed by GC-MS at the same time. The analysis was conducted in triplicates**. C)** A portion of the molecular network of GC-MS of the data collected using S’Wipe and regular sample collection approaches; an inset shows clusters of SCFAs. The pie-chart coloring for each molecule’s node corresponds to an averaged abundance of the molecule across all samples collected with the corresponding method.

### Molecular distributions captured by S’Wipe

To comprehensively assess potential method-specific biases, we compared the broader metabolome captured by S’Wipe to that obtained through conventional stool collection, examining both volatile and non-volatile molecules with GC-MS and LC-MS, correspondingly. Principal coordinate analysis shows there is a similarity of direct collection and S’Wipe, while OMNIgene Gut is significantly different. (Figure 3A. and Supplemental Table 4). We then employed molecular networking analysis(52, 53) to visualize and compare the captured molecular families between methods, which indicated that the abundances of various SCFAs are generally comparable and consistent for both methodologies (Figure 3C). The LC-MS analysis suggested that non-volatile biogenic molecules - including amino acids, bile acids, vitamins, and various microbial metabolites - were also captured similarly by both methods. The only notable differences between S’Wipe and direct collection were background molecules, primarily polyethylene glycols (PEGs) contributed by the collection matrix (Figure 3C and Supplementary Figure S4 (10.6084/m9.figshare.28404827, https://figshare.com/articles/figure/Figure_S4/28404827), S5 (10.6084/m9.figshare.28404872, https://figshare.com/articles/figure/Figure_S5/28404872)).

These findings indicate that S’Wipe collection preserves the native stool metabolome without significant alteration. After appropriate normalization and blank subtraction to account for background signals, data from S’Wipe samples should be directly comparable to direct sampling methods that do not bias metabolome composition.

### Cell debris removal

To confirm the suitability of the S’Wipe method for streamlined metabolomics workflows, we assessed its compatibility with mass spectrometry (MS) analysis without the need for extensive sample manipulation and cleanup. In MS-based metabolomics, various methodologies such as lyophilization/reconstitution, solid-phase extraction, and bi-phase extraction are commonly employed to remove unwanted background components, including proteins, lipids, and cell debris, which can interfere with MS analysis by causing clogging, introducing background noise, and affecting analyte solubility and ionization efficiency (54–56).

Fecal samples naturally contain a large number of host and microbial cells that must be removed before introduction into the MS system (56–58). Unlike direct fecal sample collection methods, the S’Wipe approach is designed to retain the fecal material within the paper matrix while allowing soluble metabolites to partition into the solvent. We hypothesized that this feature of the S’Wipe method would result in a lower cellular content in the collected samples compared to traditional fecal collection techniques. To test this hypothesis, we conducted a comparative cell counting analysis (including both native feces and supernatant) between samples obtained using the S’Wipe approach and those collected through conventional methods. Supplemental Figure S9 (10.6084/m9.figshare.28404923, https://figshare.com/articles/figure/Figure_S9_/28404923?file=52314203) is showing mass of stool collected in milligrams on the secondary axis (blue bars) as well as Normalized density (ND) of samples as determined by CellsBin proprietary technology (description is given in the Methods section) of real time cell counting (Supplementary Movies (10.6084/m9.figshare.28462592, https://figshare.com/articles/media/Supplemental_Movies_/28462592)). It is evident that the abundance of metabolites is similar between conventional stool collection and S’Wipe, as discussed above, but the cells are generally absent in S’Wipe samples, ostensibly due to their retention on the tissue. Minimizing the presence of cell debris mitigates the risk of instrumentation contamination and circumvents issues such as clogging that may arise from excessive cellular material, enabling a simplified extraction protocol, where a single centrifugation step yields a supernatant that is ready for direct MS analysis.

### S’Wipe use for DNA collection

Finally, we investigated the potential for “multiomix” sampling - dual metabolomics/microbiome analysis using S’Wipe. We collected samples from both adult and infant subjects over three time points (days 1, 3, and 5) and compared them against samples collected using DNA/RNA Shield buffer, a widely-used commercial solution optimized for microbiome preservation and DNA extraction (**Figure 4**). After centrifugation, both the supernatant and the collection paper were subjected to DNA extraction (see Methods). We found that DNA was fully retained on the paper material, with no detectable DNA in the supernatant. This is particularly advantageous for multi-omics applications, as DNA-free supernatant is optimal for metabolomics analysis, preventing column contamination and clogging, while DNA retention on the paper captures genomic material for sequencing.

**Figure 4.**
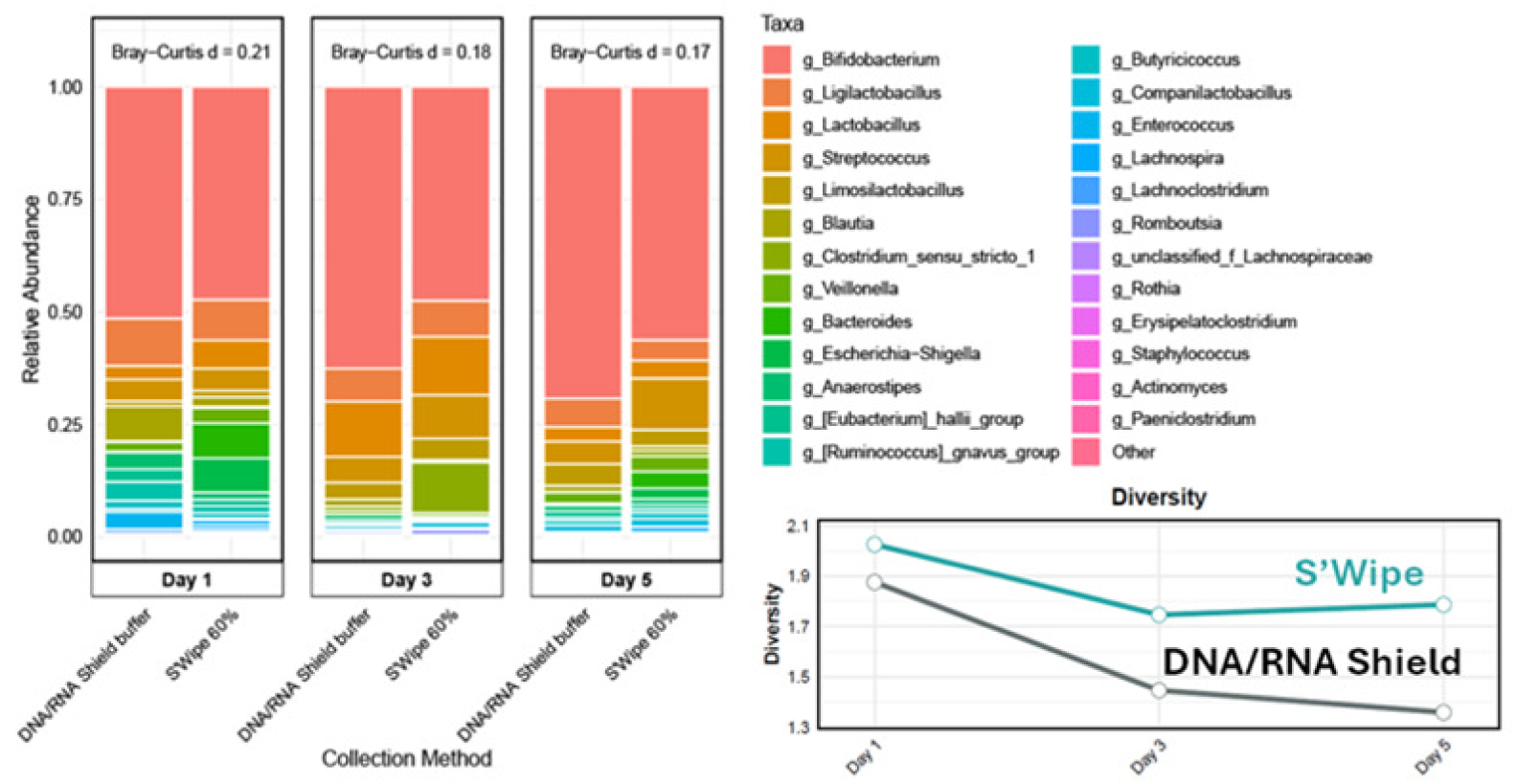
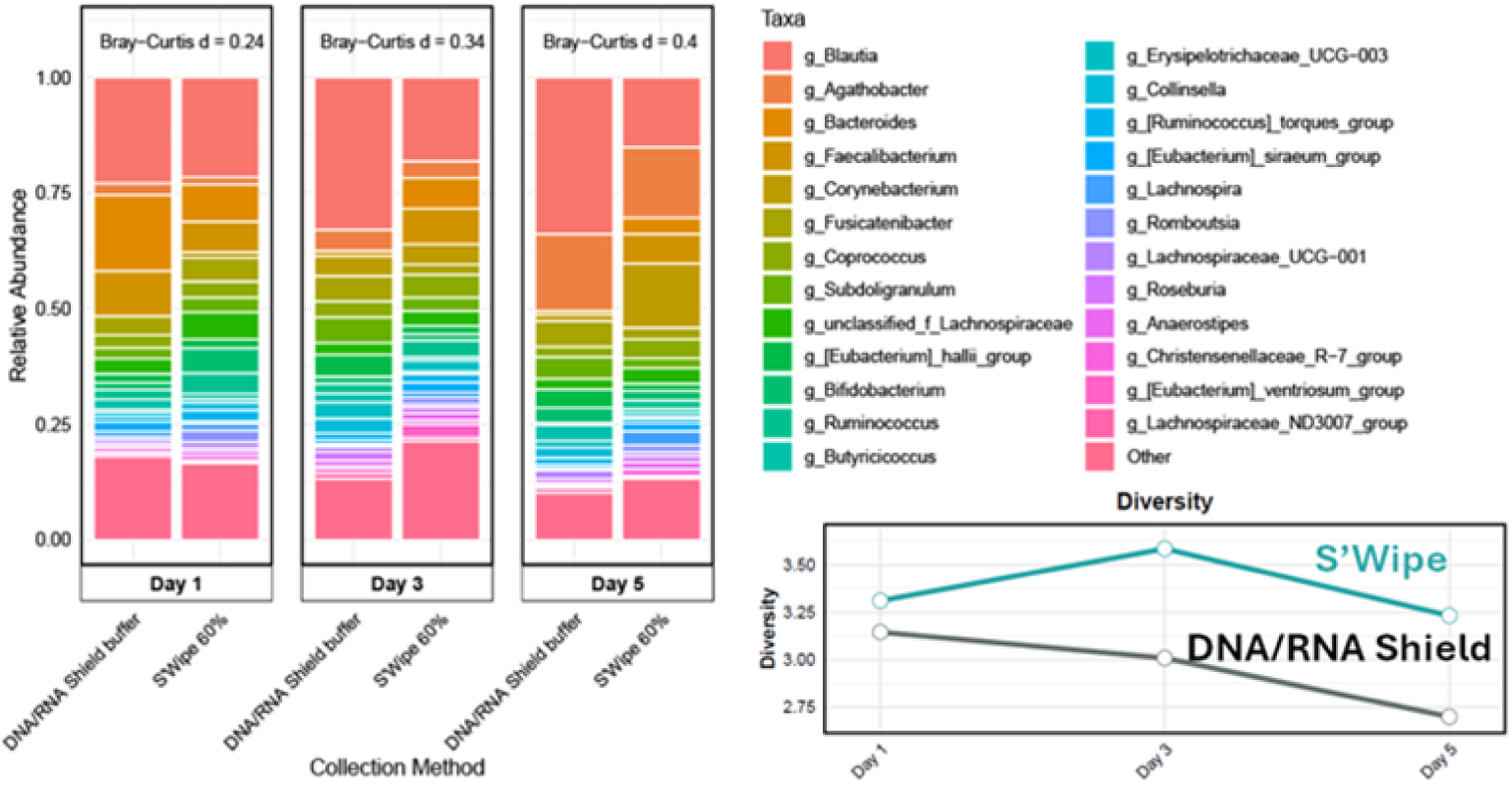
16S Top 25 Genera for DNA/RNA Shield vs. S’Wipe collection. **A)** Infant **B)** Adult. Each panel displays paired samples collected using DNA/RNA Shield buffer (left) and S’Wipe (right) at three independent collections from the same individuals (days 1, 3, and 5). Bray-Curtis dissimilarity values between methods are shown above each time point. Accompanying diversity plots (right) show temporal changes in alpha diversity for both collection methods.

16S rRNA gene sequencing revealed that S’Wipe collection does efficiently capture microbial DNA and it appears to be preserved by the ethanol buffer during storage and shipping, with no apparent bloom noticeable. Moreover, S’Wipe samples consistently showed higher microbial diversity compared to DNA/RNA Shield samples across all time points (**Figure 4**). As expected for developing gut microbiota, infant samples showed lower overall diversity compared to adult samples. Also, both groups exhibited the same trend - S’Wipe captured slightly greater microbial diversity than DNA/RNA Shield. The difference was particularly pronounced in adult samples, where S’Wipe detected several additional taxa, including members of the Erysipelotrichaceae and Christensenellaceae families. We noted the presence of some skin-associated genera such as Staphylococcus in S’Wipe samples, likely due to contact with perianal skin during collection. After excluding these skin-associated taxa, S’Wipe samples maintained higher diversity of gut-specific microorganisms compared to DNA/RNA Shield samples. The Bray-Curtis dissimilarity metrics between the two methods remained relatively stable for infant (0.17-0.21) and adult samples (0.24-0.40), suggesting consistent capture of major community structures by both sampling methods. These findings demonstrate that S’Wipe enables simultaneous collection of high-quality samples for both metabolomics and microbiome analyses, potentially simplifying multi-omics study designs.

## DISCUSSION

The gut represents one of nature’s most complex molecular environments, shaped by dietary inputs, environmental exposures, host metabolism, and microbial transformations. To develop reliable metabolite-based diagnostics, particularly for key molecules like short-chain fatty acids (SCFAs), sampling methods that can scale to capture this complexity across large, diverse populations are needed. Current approaches cannot meet the demand for collecting and analyzing thousands to millions of samples needed to routinely track clinical biomarkers. This provided the motivation for a simple, cost-effective stool collection method designed for population-scale metabolomic studies.

Sequencing advances have enabled massive initiatives like the Human Microbiome Project (59–66) and the Earth Microbiome Project (EMP) (67–69). Unlike DNA sequencing, where the target molecule is consistent, enabling joint co-analysis of collected data (59–61, 69), metabolites vary widely in their chemical properties, making standardization complex. This creates a bottleneck: the lack of standardized methods hinders large-scale studies, while the absence of studies comparable to EMP(62)(70) reduces pressure for methodological standardization.

A “universal” metabolomics sampling method must satisfy three key criteria: 1) collect native samples without introducing bias, 2) minimize distortions from sample handling and preservatives, and (3) remain compatible with various analytical platforms (GC-MS(71–73), LC-MS (74–78), and NMR (79–81)). Additionally, it must be cost-effective, user-friendly, and scalable - avoiding critical materials, particularly metals (54, 82–87). These requirements differ fundamentally from DNA sequencing, where uniform protocols can be applied across all samples.

The S’Wipe collection method was designed with simplicity and the ease of use as the primary goal, building on the most basic element of the bathroom routine - toilet paper. The simplicity also makes sampling manifold cheap (<$1 per kit) and easy to manufacture. More importantly, the approach minimizes user burden, enabling individuals to adopt it without disrupting their established routines. While S’Wipe doesn’t fundamentally alter the per-sample cost of metabolomic analysis (as sampling cost is relative minor compared to downstream MS analysis), it can enable true economies of scale by removing logistical barriers that make current collection methods impractical for population-level studies. Just as commercial testing laboratories process millions of routine clinical samples annually, metabolomics could achieve similar scale once sampling is simplified enough to be practically implemented at that level.

Using the toilet wipe allows collecting neat fecal samples without alteration, as evidenced by the results presented above (**Figure 3A** and Supplementary **Figure S4 (**10.6084/m9.figshare.28404827, https://figshare.com/articles/figure/Figure_S4/28404827)).

Since the samples collected by S’Wipe are directly comparable to neat stool samples, data from any study utilizing S’Wipe, in principle, should be directly comparable to studies where neat stool is collected by any other means, enabling continuation of existing studies, building biobanks, routine sampling, cross-laboratory data comparison, etc., provided standardization of downstream analysis protocols.

We selected 60% ethanol as a versatile preservation medium. This solvent effectively captures many diagnostically relevant metabolites, including SCFAs, while being cost-effective and relatively non-toxic. Ethanol suppresses microbial growth and enzymatic activity(88), with our data showing stable SCFA levels even after days at room temperature (**Figure 2B**). While the S’Wipe protocol uses ethanol by default, it can be adapted for specific applications - samples can be lyophilized and reconstituted in alternative solvents to optimize detection of particular molecular classes. However, when targeting molecules beyond SCFAs, their stability in ethanol should be validated to ensure reliable results. In addition to the validated analysis protocol, the S’Wipe workflow can be modified for specific applications. For example, samples can be lyophilized and reconstituted in alternative solvents to optimize detection of specific molecular classes (this approach is only suitable for non-volatile compounds). The choice of solvent can be tailored to specific analytical methods (e.g., RP vs. HILIC for LC-MS).

The amount of collected material varies naturally; sample weight can be determined by subtracting the standardized weights of the collection tube and wipe from the total kit weight. Although metabolite concentrations may poorly correlate with total mass of stool due to variations in water and fiber content, S’Wipe collection resulted in strong correlation between SCFA abundance and collected biomass, validating our mass-based normalization approach (**Figure 2A**).

We have demonstrated that S’Wipe method separates metabolites (in supernatant) from cells and DNA (retained on cellulose), with no cell debris detected in the supernatant (Supplemental Figure S9 (10.6084/m9.figshare.28404923, https://figshare.com/articles/figure/Figure_S9_/28404923?file=52314203)). This separation streamlines sample processing and allows both metabolomics and microbiome analysis from a single collection event - an advantage over traditional approaches that require separate sampling protocols for each “omics” measurement.

This dual-purpose capability can benefit integrative studies by reducing batch effects and simplifying sample pairing between metabolomic and microbial profiles. The approach is particularly useful for longitudinal intervention studies where tracking both microbial communities and their metabolic outputs is needed. By preserving stool metabolites and microbial DNA while reducing sample cleanup requirements, S’Wipe provides a straightforward, cost-effective option for large-scale gut health assessments. The method’s ability to support native sample collection while performing comparably to specialized collection methods provides a new tool for population-wide multi-omics studies.

## Conclusion

This study describes a fecal sampling routine designed for high-frequency metabolomics sampling. The approach condenses into the ubiquitous toilet wipe use without compromising analytical sensitivity or quantitative molecular preservation. We have shown the intrinsic capacity to stably sorb, store, and release key gut metabolites such as SCFAs and bile acids using a simple paper matrix, suitable for self-administration. By simplifying the stool collection process, this approach is designed to translate monitoring of the gut microbiome on the molecular level from an academic, into a practical consumer tool for personalized nutrition and disease management. Similar to how affordable genetic testing has allowed proactive mitigation of hereditary risks, routine measurement of molecular biomarkers such as SCFAs is needed to shift gastrointestinal disease management from reactive to predictive by providing easy access to gut metabolome monitoring.

## Materials and Methods

### S’Wipe kits

All of the conducted studies have utilized S’Wipe kits manufactured by Arome Science Inc. The kits contained lint-free paper (Kimberly-Clark Professional™, 34120), the preservation solution in a 5 ml container spiked with an internal standard, a tube stand, and a sampling instruction sheet. The kits were provided to study participants free of charge.

### Protocol Optimization Study

A healthy volunteer donor has provided all stool samples. The protocol described in the “Sample Extraction” section was run alongside a long sonication and incubation extraction protocol. The long protocol included a 10 minute sonication on ice and six hours on a vibration table at 2 - 4°C for homogenization. Samples included S’Wipe kits with no wipe, S’Wipe kits complete, and S’Wipe kits with no stool. Samples were generated in triplicates. After the sample extraction, samples were analyzed by GC-MS and LC-MS/MS.

### Stability Study

A healthy volunteer donor has provided all stool samples. Samples collected by S’Wipe kits and controls were incubated at 25°C for 0-, 3-, 5-, and 7-days. Negative controls contained no stool, and positive controls contained no stool and Accustandard FAMQ-004 at 1mM. All samples were collected in triplicates. Sampled kits were stored at −80°C until analysis and all samples were extracted together. After the sample extraction, samples were analyzed by GC-MS and LC-MS/MS.

### Shipping Study

A healthy volunteer donor has provided all stool samples. Samples collected by S’Wipe kits and controls were sent and returned via United States Postal Service to a short-distance (36-miles) and long-distance (2,950-miles) address. Negative control contained no stool. All samples were in triplicates. Upon reception in the mail, the S’Wipe kits were stored at 2 - 4°C until analysis. All samples were extracted and analyzed by GC-MS and LC-MS/MS at the same time.

### Interpersonal Variability Study

10 healthy volunteers were provided with S’Wipe collection kits and US mail return labels to return samples to the lab. No personal or any other information was collected throughout the study.

### Sample Extraction

Different parameters such as the order of steps, solvent volume, and centrifugation time and setting were tested to establish the standard protocol, which was selected as the simplest and most effective approach. Completed S’Wipe kits kept at −80°C were thawed on ice prior to extraction. Pre-sample and post-sample kit mass was recorded. The completed kits were first placed in an iced ultrasonic bath for 10 minutes and then on a vibration table for 10 minutes at room temperature for homogenization. 1000μL aliquot was taken and transferred to 1.5 mL microcentrifuge tubes. The samples were centrifuged for 5 minutes at 14,000 revolutions per minute. After centrifugation, aliquots of 100 uL were taken from each sample and transferred to vials with conical inserts, and analyzed by GC-MS and/or LC-MS/MS. Samples were directly loaded onto GC-MS. For LC-MS/MS, samples were diluted 4-fold with pure cold methanol to precipitate protein. The samples then were filtered through Phenomenex Phree plates with the application of 4 psi negative pressure. 200 uL of collected filtrates were dried under vacuum and reconstituted with 100uL of C18 resuspension buffer (5% Acetonitrile (Sigma, USA) in LCMS grade water with added internal standards: Sulfachloropyridazine (TCI, USA) and Sulfamethazine (Sigma, USA).

### GC-MS data acquisition and processing

1 μL aliquot of the S’Wipe kit supernatant was directly injected into an Agilent 6890 GC interfaced to a mass spectrometer Hewlett Packard MSD 5973 for electron ionization GC-MS. The GC utilizes a 30m ZB-FFAP column (0.25 mm i.d., 0.25 μm film thickness) for metabolite separation with 1.2mL/min constant He flow. The oven temperature program initiates at 50°C rising to 240°C at 10°C/min. No noticeable carryover was observed over the entire injection sequence for all of the studies. Also, no increased contamination that necessitates liner change has been observed, indicating that possible lint particulate traces from wipe material did not contribute no observable analytical interference. The data were then deconvoluted with the MSHub algorithm (52). The experimental spectra were searched against the NIST 2023 library with ≥80% spectral match defining putative identifications, and the retention times falling within <0.01 min of the corresponding reference standards. Targeted analysis of SCFAs and p-cresol was performed by using Agilent Masshunter software.

### LC-MS/MS data acquisition and processing

The samples were injected and chromatographically separated using a Vanquish UPLC (Thermo Fisher Scientific, Waltham, MA), on a 100 mm × 2.1 mm Kinetex 1.7 μM, C18, 100 Å chromatography column (Phenomenex, Torrance, CA), 40 °C column temperature, 0.4 mL/min flow rate, mobile phase A 99.9% water (J.T. Baker, LC–MS grade), 0.1% formic acid (Thermo Fisher Scientific, Optima LC/MS), mobile phase B 99.9% acetonitrile (J.T. Baker, LC–MS grade), 0.1% formic acid (Fisher Scientific, Optima LC–MS), with a the following gradient: 0–1 min 5% B, 1–8 min 100% B, 8–10.9 min 100% B, 10.9–11 min 5% B, 11–12 min 5% B.

MS analysis was performed on Orbitrap Exploris 240(Thermo Fisher Scientific, Waltham, MA) mass spectrometer equipped with HESI-II probe sources. The following probe settings were used for both MS for flow aspiration and ionization: spray voltage of 3500 V, sheath gas (N_2_) pressure of 35 psi, auxiliary gas pressure (N_2_) of 10 psi, ion source temperature of 350 °C, S-lens RF level of 50 Hz and auxiliary gas heater temperature at 400 °C.

Spectra were acquired in positive ion mode over a mass range of 100–1500 *m*/*z*. An external calibration with Pierce LTQ Velos ESI positive ion calibration solution (Thermo Fisher Scientific, Waltham, MA) was performed prior to data acquisition with ppm error of less than 1. Data were recorded with data-dependent MS/MS acquisition mode. Full scan at MS1 level was performed with 30K resolution. MS2 scans were performed at 11250 resolution with max IT time of 60 ms in profile mode. MS/MS precursor selection windows were set to *m*/*z* 2.0 with *m*/*z* 0.5 offset. MS/MS active exclusion parameter was set to 5.0 s.

LC-MS raw data files were converted to mzML format using msConvert (ProteoWizard). MS1 features were selected for all LC-MS data sets collected using the open-source software MZmine 3 (89) with the following parameters: mass detection noise level was 10,000 counts, chromatograms were built over a 0.01-min minimum time span, with 5,000-count minimum peak height and 5-ppm mass tolerance, features were deisotoped and aligned with 10-ppm tolerance and 0.1-min retention time tolerance, and aligned features were filtered based on a minimum 3-peak presence in samples and based on containing at least 2 isotopes. Subsequent blank filtering, total ion current, and internal standard quality control were performed.

### Untargeted metabolomics data processing

MetaboAnalystR (Chong and Xia, 2018) was used to preprocess the data by replacing zero peak abundance values with 1/5 of the smallest positive value for each feature, representing the limit of detection. Peak abundances below 1 (4.4% of data) were adjusted to 1 to prevent negative values during log transformation. This preprocessing step ensures compatibility with downstream batch correction tools (e.g., ComBat, WaveICA2) and avoids issues to negative values in the corrected matrix. Within the overall data distribution, metabolite features with such low peak abundances (<1) are generally biologically insignificant and likely stem from zeros in the raw data. Features with no variance (constant or single value across samples) were removed. Quantile normalization was performed to standardize distributions across samples to reduce technical variability, followed by centered log-ratio (CLR) transformation. PERMANOVA, implemented using the adonis2 function from the “vegan” package in R (number of permutations = 999), was performed to test whether the multivariate means of S-wipe and stool samples differed significantly, using the Euclidean distance matrix of the CLR-transformed feature table as input.

### Cell counting assay

Blind samples were analyzed using CellsBin’s VEGA platform at an external lab. The normalized density of each sample was calculated by imaging 20 µl through CellsBin’s microfluidic optical device. The company’s machine learning algorithm detected and classified particles, estimating the relative size and normalized density of each sample.

### Stool sample collection, DNA Extraction and 16s rRNA V4 region sequencing

Stool samples were initially collected on tissue wipes, which were carefully folded to prevent contamination and ensure the stool remained securely on the wipe’s surface. These tissue wipes were then transferred into 50 mL Falcon tubes pre-filled with 60% of ethanol or DNA/RNA Shield. A blank control tube was also included, containing DNA/RNA Shield but no wipe with stool sample, to monitor potential contamination from the tissue wipe or reagents. All samples were stored at 4°C. To extract DNA from the wipe samples, the tissue wipe remained in the Falcon tube, and a 1 mL pipette tip was used to press the wipe repeatedly to extract the liquid absorbed within it. This process yielded approximately 100–300 µL of liquid per sample. The extracted liquid was then centrifuged at 15,000g for 1 minute, resulting in the formation of a visible stool pellet at the bottom of the tube. This pellet was subsequently used for DNA extraction using the Qiagen PowerSoil DNeasy Pro Kit. The V4 region of 16S rRNA was amplified and sequenced at the Illumina Miseq platform.

## Data availability

All data generated in this study are publicly available. The raw data are available on MassIVE Repository (massive.ucsd.edu) under the following dataset accession numbers [MSV000094530](https://massive.ucsd.edu/ProteoSAFe/dataset.jsp?task=7942891df5cf4807a4c7751256dc5a66) (GC-MS), [MSV000094529](https://massive.ucsd.edu/ProteoSAFe/dataset.jsp?task=d878619cb817458f9e118c1607ea6758)(LC-MS), and [MSV000096969](https://massive.ucsd.edu/ProteoSAFe/dataset.jsp?task=bc328ae3a59c45218c8a128d4b327caa) Reproducibility data (GC-MS). https://gnps.ucsd.edu/ProteoSAFe/status.jsp?task=eab1df9e88584bf289969bceb0ca59a3 https://gnps.ucsd.edu/ProteoSAFe/status.jsp?task=bf75f44dfab149248e66fafa6dd74e0a

## AUTHOR CONTRIBUTIONS

AM and AA formulated the study. AM, KP, HM prepared the samples. AM devised the sampling manifold. AM, KP, AL performed the GC-MS and LC-MS analysis. EK, AM, DM and AL conducted data processing and statistical analysis. AA, AM, HM coordinated the study. AA, DM, AL, AM wrote the manuscript. All authors read and approved the manuscript.

## Acknowledgments

The authors appreciate the help and materials in the form of automation support provided by TomTec Inc. (www.tomtec.com). Additionally, the authors thank CellsBin Inc. (www.cellsbin.com) for the analysis of samples for microscopy measurements (CellsBin, based in New Haven, CT, is a tech-bio company. Its VEGA platform is utilized in precision oncology and other fields, including aerosol detection for the prestigious IARPA’s Picard Project (https://www.iarpa.gov/research-programs/picard)). Arome Science Inc. provided funding for the study

